# Nonlinear calcium ion waves along actin filaments control active hair–bundle motility

**DOI:** 10.1101/292292

**Authors:** J. A.Tuszynski, M. V. Sataric, D. L. Sekulic, B. M. Sataric, S. Zdravkovic

## Abstract

Actin filaments are highly dynamic semiflexible cellular biopolymers with diverse functions, such as cell motility. They also play the role of conduits for propagation of calcium ion waves. In this paper, we propose a new biophysical model that describes how actin filaments with their polyelectrolyte properties serve as pathways for calcium ion flows in hair cells. We show this can be utilized for the tuning of force–generating myosin motors. In this model, we unify the calcium nonlinear dynamics involved in the control of the myosin adaptation motors with mechanical displacements of hair– bundles. The model shows that the characteristic time scales fit reasonably well with the available experimental data for spontaneous oscillations in the inner ear. This model offers promises to fill a gap in our understanding of the role of calcium ion nonlinear dynamics in the regulation of processes in the auditory cells of the inner ear.

## 1. Introduction

In this paper we present a new concept aimed at explaining how the polyelectrolyte properties of actin filaments within the stereocilia of the inner ear allow them to become pathways for flow of Ca^2+^ ions. This behaviour is implicated in the adaptation of myosin motors making it a faster and more efficient mechanism for controlling Ca^2+^ ion movement when compared with simple diffusion processes. Following the introduction of the model, we apply it to a quantitative analysis of the detachment dynamics of myosin motors. We then establish a relationship between motor adaptation and spontaneous oscillations of hair–bundles.

Neighbouring stereocilia in hair–bundles of the inner ear are connected by tip links, which are joined to transduction channels. The deflection of the stereocilia by sound waves changes tip link tension and brings about opening and closing of transduction channels permitting or preventing the entrance of K^+^ and Ca^2+^ ions into stereciolia, respectively. The role of Ca^2+^ ions is to detach myosin motors in order to ease an increased tip link tension. Conversely, if the tension is reduced the myosin motors reattach and ascend along actin filaments restoring the resting tension in tip links. In this paper, attention is placed on the ability of hair–bundles in the bullfrog’s sacculus to produce oscillations that might underlie spontaneous otoacoustic emissions.

The paper is organized as follows. In Section 1, we explain the polyelectrolyte character of actin filaments within stereocilia. Our biophysical model that describes the mechanism of localized Ca^2+^ ion pulse propagation along actin filaments is presented in Section 2. Further, in Section 3 we discuss in detail the Ca^2+^ ion dependent myosin–based adaptation of hair–bundle motility. Finally, our model’s application to the spontaneous oscillations of hair bundles is presented and discussed in Section 4. In Section 5 we present main conclusions and discuss the obtained results.

## 2. Actin filaments as polyelectrolyte guidelines for Ca^2+^ ions within stereocilia

We first explain how actin filaments within the hair–bundles can serve as conduits for Ca^2+^ ion flows, which are implicated in myosin motor adaptation. Recently, in a separate paper we have described why and how actin filaments can be considered as linear polyelectrolytes [1], which is in accordance with the approach due to Manning [2]. It is well known that the globular actin proteins have a diameter *r* = 5.4 nm and spontaneously polymerize to form double stranded filaments referred to F–actin (see Fig. 1).

**Fig. 1.**
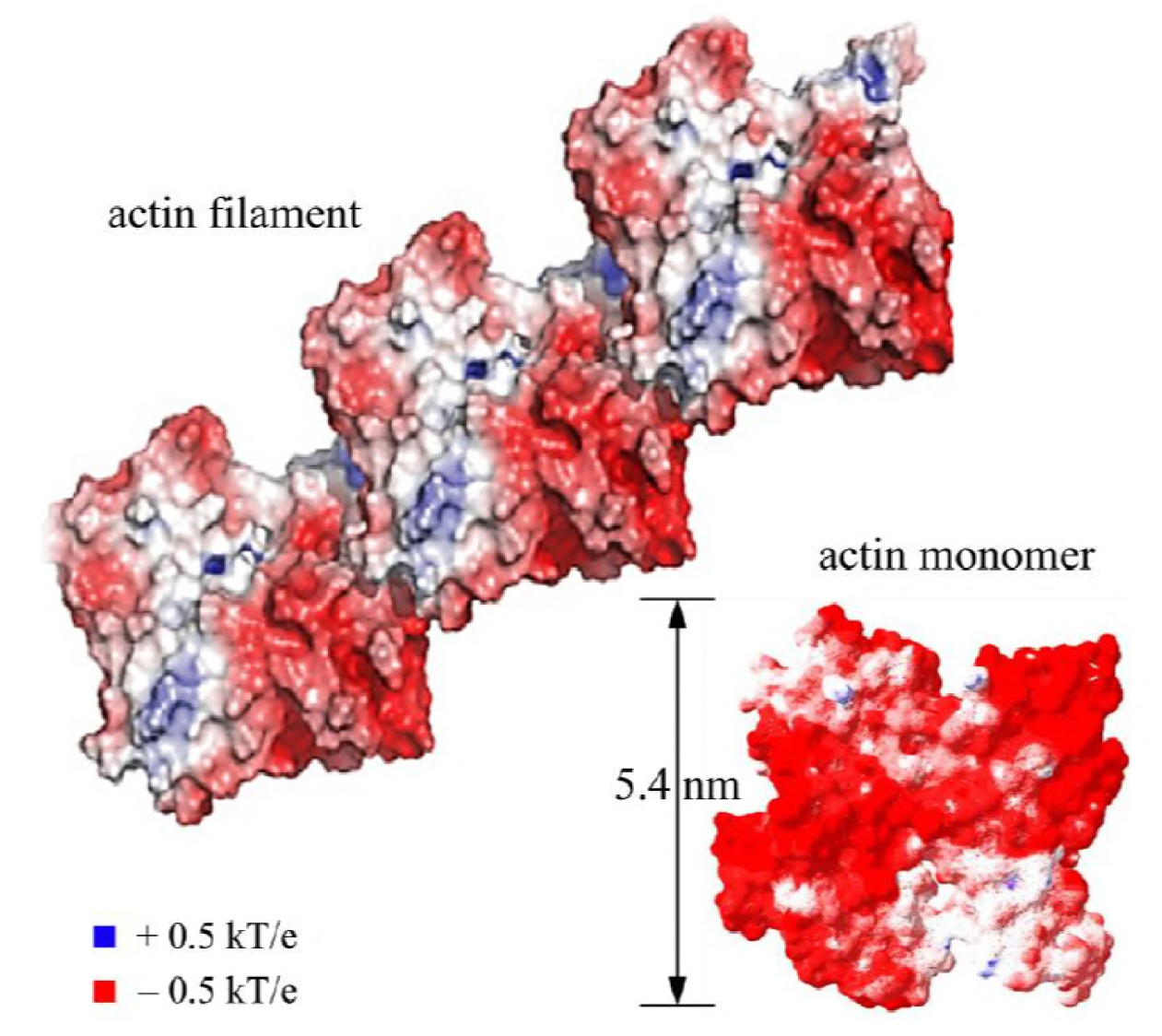
Structural model of an actin filament consisting of two intertwined long–pitch right–handed helices showing an electrostatic charge distribution on the surface of an actin monomer and a segment of the actin filament (red color indicates negative charges, blue positive charges and white neutral regions).

The polyelectrolyte nature of actin filaments at physiological conditions was discovered some twenty years ago and the crucial parameters were presented by Tang and Janmey [3]. The various theoretical and experimental results available today provide an explanation why actin filaments are true polyelectrolytes, which play functional roles in the signalling pathways for Ca^2+^ ions within the cells [4]. Our approach strongly relies on this concept.

The basic idea is based on the fact that the negatively charged surface of an actin filament attracts positive counterions from the cytosol, mainly in terms of divalent ions such as Ca^2+^. Each actin monomer carries an excess of 11 negative elementary charges *e* = 1.6×10^−19^ C, and counting two strands with two monomers one easily finds that the linear charge density is approximately 4 *e*/nm. This gives the linear spacing of elementary charges along the filament axis to be *δ* = 0.25 nm. This should be compared with a characteristic Bjerrum length *l*_B_, which defines the distance at which thermal fluctuations are equal to the electrostatic attraction or repulsion between ions and negative sites on the actin filament. For divalent Ca^2+^ ions (*z* = 2) at *T* = 310 K, the Bjerrum length is determined by

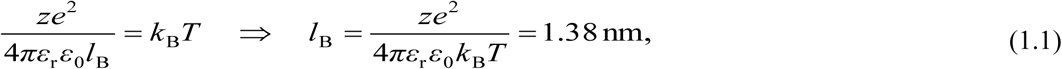

where *ε*_r_ = 78, *ε*_0_ = 8.85×10^−12^ F/m and *k*_B_ = 1.38×10^−23^ J/K is the Boltzmann constant. In accordance with this value, the corresponding dimensionless parameter *ξ* for actin filament is found as

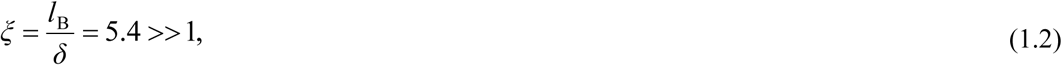

which is a very important factor for electrolytic free energy of Ca^2+^ ions around an actin filament. Since this value is significantly greater than unity, the necessary condition for the formation of a polyelectrolyte is clearly satisfied supporting our argument about the polyelectrolyte character of actin filaments at physiological conditions. This also confirms that the attractive Coulomb force brings Ca^2+^ ions very close to the filament surface forming a cylindrical condensed “ionic cloud” with a thickness on the order of *d* = 1 nm and an inner radius of approximately *r* = 5.4 nm (see Fig. 2). Around this “ionic cloud” there is a layer depleted of ions of both signs with a thickness equal to the Bjerrum length *l*_B_. Consequently, an actin filament can be considered as a transmission line with a nano–capacitor that is capable of storing and transporting a cloud of attracted Ca^2+^ ions along the length of the actin filament [5].

**Fig. 2.** Schematic representation of the counterion charge distribution surrounding an actin filament.

In order to properly apply the theory of polyelectrolytes [6], the Debye screening length Δ in the hair cell’s endolymph must first be estimated. Since the average ionic concentration in a typical endolymph is on the order of *n_s_* = 9×10^25^ m^−3^, the Debye length amounts to [7]

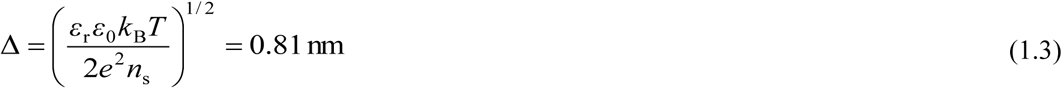

An electrolytic free energy *G* of an actin filament with *N* condensed Ca^2+^ counterions is defined as follows [6]

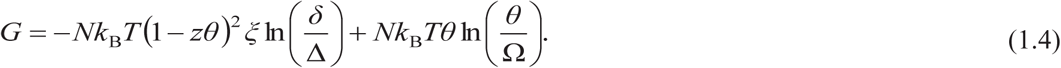

The first term in the above expression reflects the Debye–screened repulsion between all pairs of negative sites on the charged filament considered. With the already determined parameters Δ = 0.81 nm and *δ* = 0.25 nm, the first term in Eq. (1.4) is positive as required because ln(*δ*/Δ) = –1.176. The dimensionless quantity *θ* represents the relative concentration of Ca^2+^ with respect to the negative sites of the filament. If every negative site is covered by one counter ion, then the total coverage is *θ* = 1. Under a lower ionic concentration around the filament, the condition *θ* < 1 holds and it leads to the inequality

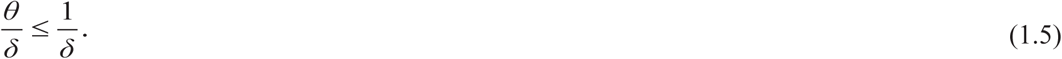

For this reason, the effective charge of a site is reduced to the following fraction for Ca^2+^ counterions

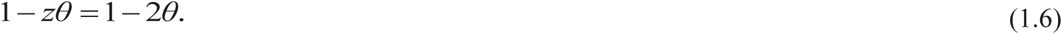

The second term in Eq. (1.4) involves an internal partition function Ω, which reflects the short–range interactions between Ca^2+^ counterions and an actin filament in accordance with the concept of an ideal gas of particles.

By using the free energy given in Eq. (1.4) we define the free energy per molecule expressed in units of thermal energy as

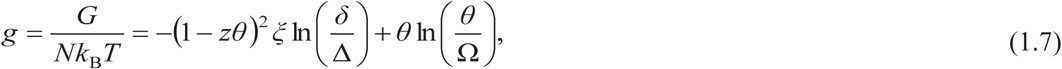

It is then possible to calculate the electrochemical potential *μ_c_* for a condensed “ionic cloud” as follows:

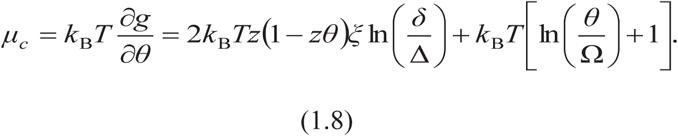

The equilibrium concentration of counterions on an actin filament can be adequately estimated by equalizing the first term of the above-defined chemical potential *μ*_c_ with the potential of free counterions *μ*_free_ for a stationary endolymph concentration (*n*_s_ = 9×10^25^ m^−3^ = 0.15 mol/l). Considering that

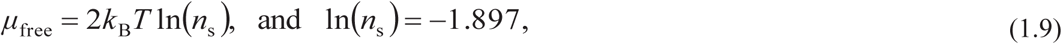

the equality will hold if we balance the multipliers of 2*k*_B_*T* in Eqs. (1.8) and (1.9)

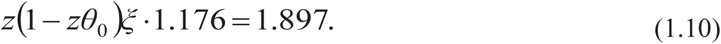

This resultant equality determines the equilibrium concentration of a partial coverage of an actin filament by Ca^2+^ counterions as

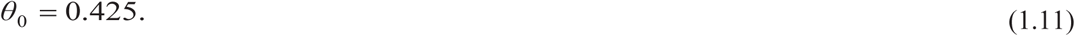

The fraction of remaining negatively charged sites on an actin filament is equivalently expressed as:

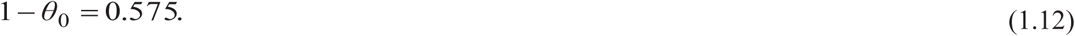

This number defines the space such that with an increased influx of Ca^2+^ ions through transduction channels, the additional ions can be accommodated on the filament creating a denser “ionic cloud”. The ions from the “ionic cloud” are free to slide along the nearby actin filament but are inhibited from diffusing away. This delocalized binding is more preferable for counterions of higher valence such as Ca^2+^ ions, in which case the charged filament is neutralized to a higher degree.

Let us estimate the concentration of Ca^2+^ ions within an “ionic cloud” under thrmodynamic equilibrium conditions. Using a well–known fact that one elementary unit of an actin filament has 2×11*e* = 22*e* charges with an estimated fractional covering of *θ*_0_ = 0.425, so the number of Ca^2+^ ions is approximately n = 9. Taking dimensions from Fig. 2, we obtain the ionic density within the thin hollow cylinder with base area A and length L = 5.4 nm as

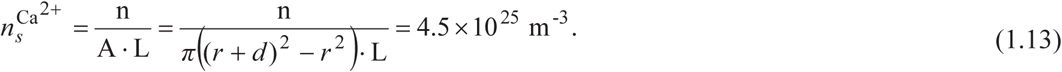

This is very close to an equilibrium ionic concentration in an endolymph, namely *n*_s_ = 9×10^25^ m^−3^. It is obvious that with a doubled number of Ca^2+^ ions within an “ionic cloud”, we find 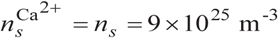, a number which is close to the global ionic concentration in an endolymph. Why is this result so important? Despite the fact that Ca^2+^ ions play a crucial role in controlling myosin motors in stereocilia, their concentration is marginally small in an endolymph compared with those of the dominant K^+^ and Na^+^ ions. The experimental assay by Beurg *et al*. [8] revealed that when the influx current through channels of a hair–bundle is 3.4 nA, just 0.2 % of this current is carried by Ca^2+^ ions with a corresponding peak of 7 pA. Moreover, some experimental data show that the total concentration of neutral Ca atoms in these processes is by one order of magnitude higher than its ionized fraction. Moreover, Gartzke and Lange [4] pointed out that the bundles of actin filaments in stereocilia form strong diffusion barriers for ionic propagation. Additionally, the presence of fixed and mobile buffers and extruding pumps also diminishes the number of diffusing Ca^2+^ ions in an endolymph, thus lowering their affordability for myosin control. Taking into account all these observations, we are convinced that this newly proposed mechanism of polyelectrolyte pathways can describe how to efficiently distribute Ca^2+^ ions along actin filaments in order to provide highly concerted hair cell’s response to fast and complex auditory signals.

Over the last two decades or so, signalling along actin filaments has been studied using both experimental and theoretical approaches. The experimental results in the early stages [9] revealed how the ionic currents in the presence of actin filaments were amplified in comparison with actin-free solution. Accordingly we earlier offered a biophysical model based on the polyelectrolyte properties of actin filaments, which emulates the nonlinear electric transmission line [5]. Below, we briefly discuss the main estimates arising from that model. Since an “ionic cloud” can be approximated by a cylindrical layer shown in Fig. 2, we could first assess the elementary capacitance of one actin dimer of length L = 5.4 nm with radius *r* = 5.4 nm and the distance between “charged plates” assumed to be *l*_B_ = 1.38 nm as follows

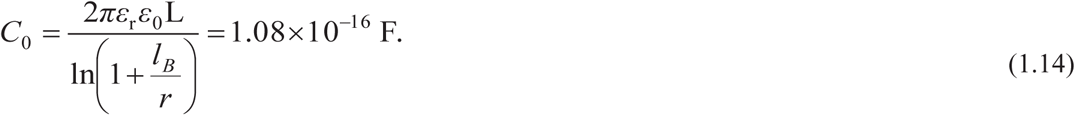

Otherwise the resistance of this same electric elementary unit of the “transmission line” can be calculated using the formula

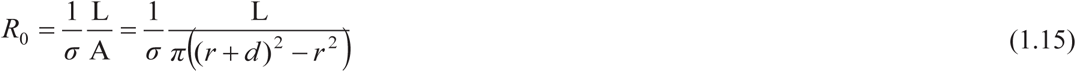

If we use two different values for the electrical conductivity experimentally determined by Minoura and Muto [10] σ^MM^ = 0.15 S/m and by Uppalapati *et al*. [11] σ^U^ = 0.25 S/m, respectively, we obtain two estimates for the elementary resistance of an “ionic cloud”, namly

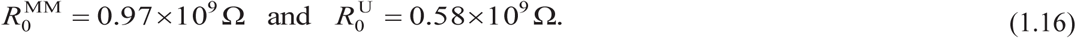

Based on the above results, we determine the characteristic times of discharging an elementary electric unit of an actin filament

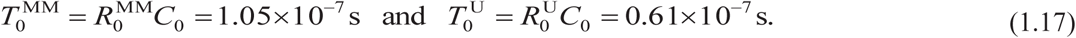

As a result, the order of magnitude estimate for the velocity of an “ionic cloud” propagating along an actin filament is

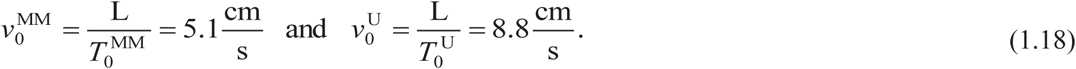

A similar order of magnitude estimate was obtained in a very recent experimental study of ionic conductivity along microtubules [12].

## 3. Equation of motion for the propagation of Ca^2+^ ions along actin filament

Although the magnitude of the velocity of “ionic cloud” propagation may be overestimated due to the use of a simplified geometry, it most likely is much greater than the velocity of diffusion of ions in bulk endolymph which is on the order of μm/s. This is comparison is our motivation for introducing the concept of fast and efficient control of adaptation myosin motors involved in the mechanoelectrical transduction in stereocilia, which we postulate is performed by the action of Ca^2+^ “ionic clouds” propagating along actin filaments.

The influx of Ca^2+^ ions through a transduction channel is mainly distributed along closely distributed actin filaments in the form of “ionic clouds”. This particular influx involves a voltage gradient in the form of a localized field *E*, which may depend on space and time. Accordingly, the local concentration of Ca^2+^ ions within an “ionic cloud” becomes a function of both time and position *x* along the actin filament. The excess Ca^2+^ ions due to channel influx propagate along an actin filament with drift velocity *v_d_* provided by the force *F* defined with a negative gradient of the electrochemical and electrical potential as follows

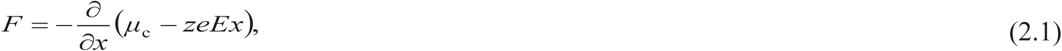

where the electrochemical potential *μ_c_* for the condensed “ionic cloud” satisfies Eq. (1.8). This force acting on Ca^2+^ ions is balanced by the viscosity damping characterized by the parameter *λ* and drift velocity *v_d_*

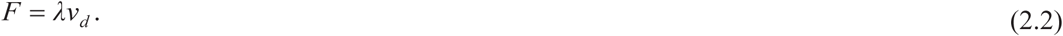

The ionic flux can be expressed as

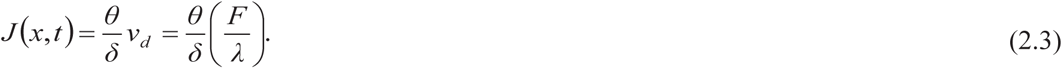

Combining the expressions given in Eqs. (1.8), (2.1) and (2.2) one finds

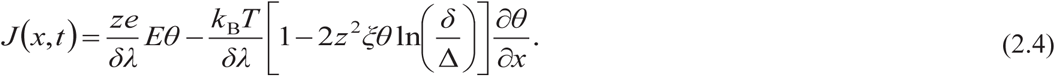

On the other hand, the linear concentration of condensed Ca^2+^ counterions must obey the continuity condition

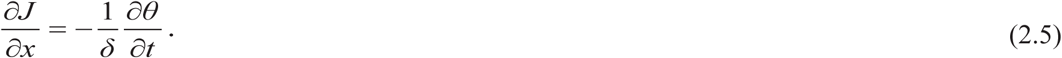

Thus on the basis of Eqs. (2.4) and (2.5) the master equation of motion for the concentration of counterions within a mobile “ionic cloud” has the following shape:

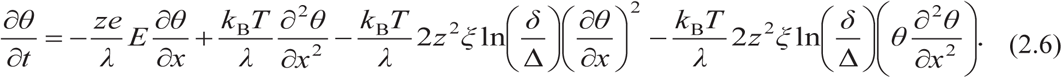

Instead of linearizing the above master equation as Manning did earlier [6], we have exactly solved this problem by using a traveling wave approach, which is widely exploited in the theory of soliton waves [13]. This makes sense keeping in mind that in Eq. (2.6) the dispersive term 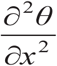 competes with the nonlinear terms 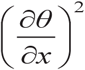 and 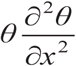 resulting in a form of balance, which leads to a moving form of the underlying “ionic pulse”.

Using a detailed derivation presented in Appendix 1, we have obtained the following implicit function *θ*(*φ*):

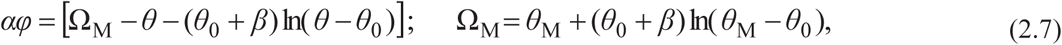

where the parameters are determined by the following expressions

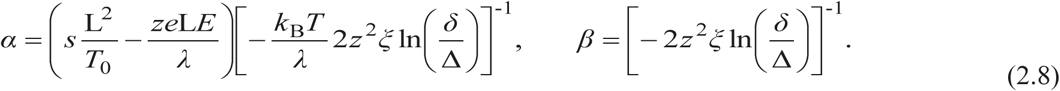

Additionally, *θ*_M_ represents the local maximum concentrations of Ca^2+^ along an actin filament, while *θ*_0_ has the value given in Eq. (1.11). Furthermore, the total coverage of negative charges of an actin filament with Ca^2+^ ions could lead to *θ*_T_ = 1 which gives the maximal ratio

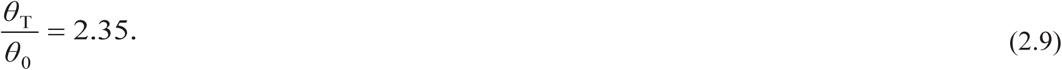

Finally, we estimate the parameters of Eq. (2.7) in order to obtain a final equation for numerical analysis. It is very easy to calculate parameter *β* on the basis of its definition given by Eq. (2.8) and the previously obtained parameters involved therein. Hence

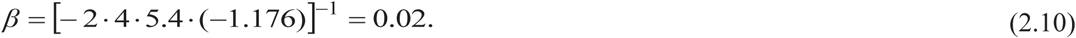

In order to calculate the parameter *α* from Eq. (2.8), it is necessary to first estimate the ratio

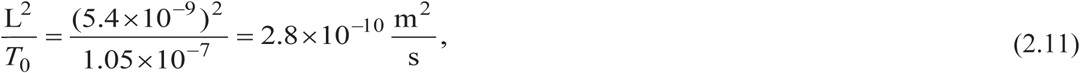

which has the unit of a diffusion constant. In an attempt to see how the applied field *E* affects the above propagation process, the viscous damping coefficient *λ* should be specified. Viscosity acting on a single Ca^2+^ ion can be roughly estimated by using the well–known Stokes’ law

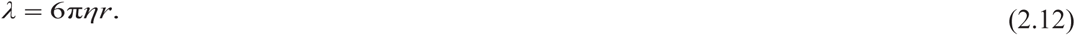

Taking water as solvent, its viscosity coefficient is *η* ~ 10^−3^ Pas. Using this value with an effective radius of Ca^2+^ ions of *r* = 2×10^−10^ m, one obtains λ = 3.7×10^−12^ Ns/m. This is a very approximate underestimation since Dhont and Kang [14] revealed that this parameter should be 20 times greater according to an experiment with Ca^2+^ ions moving around an fd virus. We will accept their estimation, which amounts to 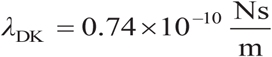.

By using both of the above parameters, the second term in Eq. (2.8) is on the order of

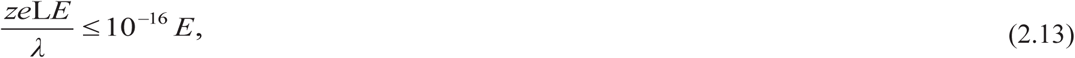

and can compete with the first one for very strong fields of the order of 10^6^ V/m. In fact, the transmembrane field is known to reach this order of magnitude [15], but we here disregard such cases using just the first term in Eq. (2.8) thus estimating the value of parameter *α* as

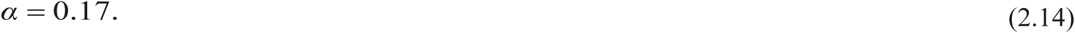

As an illustration, if we take *θ*_M_ = 2*θ*_0_= 0.85 we find that

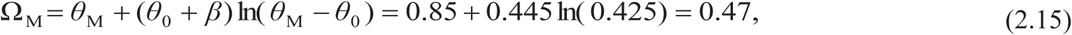

and accordingly the explicit function for Ca^2+^ ions distribution within an “ionic cloud” has the following form

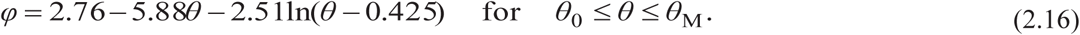

If *θ*_M_ = 0.7 the function given by Eq. (2.7) reads

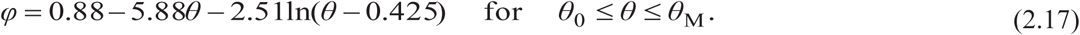

The graphical shape of the above function representing the ionic pulse is shown in Fig. 3.

**Fig. 3.**
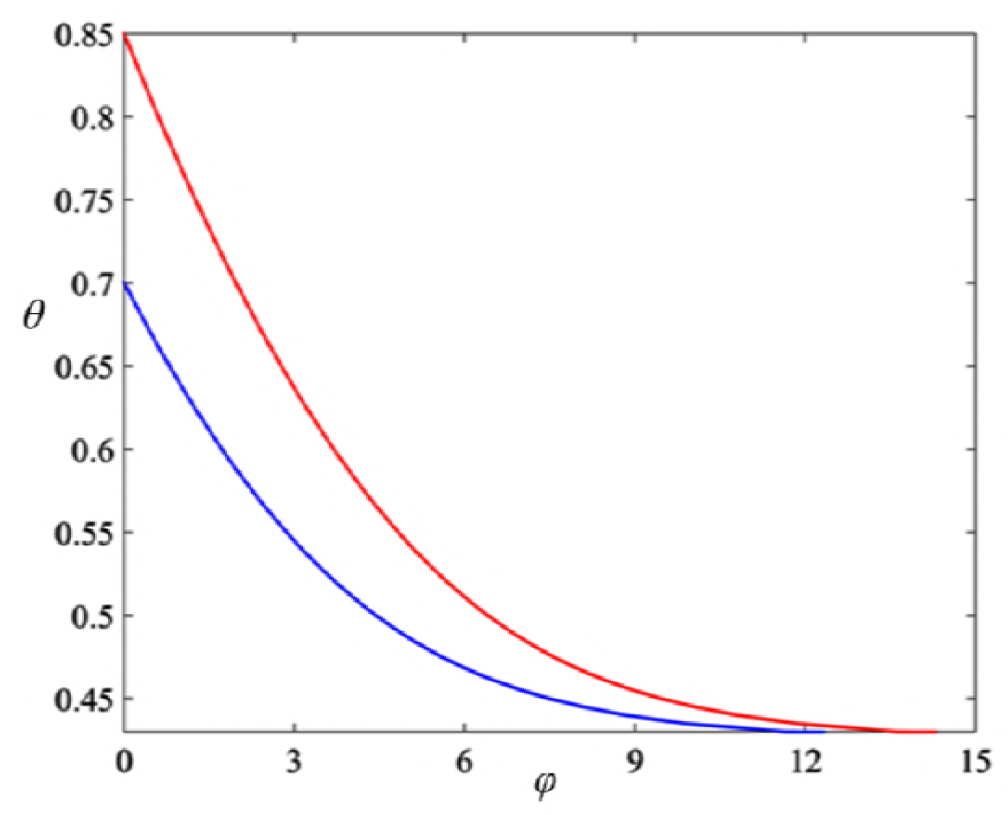
The numerical representation of functions of the model for charge distribution of Ca^2+^ counterions within a localized pulse, which propagates along an actin filament.

## 4. Ca^2+^ dependent myosin based adaptation of hair–bundle motility

In hair cells of the inner ear mechanical stimulation supplies the energy sufficient to open transduction channels responsible for the influx of positive ions. When an incoming sound wave excites the cochlea’s basilar membrane into resonant oscillation, the tectorial membrane impairs force to the mechano–sensitive hair–bundle [16]. Each hair cell is adorned with its hair–bundles, which comprise hexagonal arrays of so-called stereocilia, and play the role of mechanosensory antenna and force generator for amplificatory processes. Every stereocilium is a bundle of parallel actin filaments interconnected with some specific lateral links. The tip links are of basic importance since they serve for opening and closing of ionic transduction channels of the hair cell membrane. The hair–bundle is arranged like a harp or the back acoustic compartment of a piano in rows of increasing height (see Fig. 4). As an illustrative example, in a striolar hair–bundle of chicks the longest stereocilia are 9.2 μm and the shortest are 2.4 μm. The number of stereocilia in that bundle is N = 51. The diameter of a single stereocilium is about 0.42 μm and the number of tip links in this bundle is 42. Generally, if a hair– bundle comprises N stereocilia, which slope up against each other and the longest of them has the length 𝓛, Holt and Corey [17] noted that there is the inverse relation between N and 𝓛 in a way that N spans 20–300 stereocilia whose lengths 𝓛 are in the inverse range 30–2 μm.

**Fig. 4.**
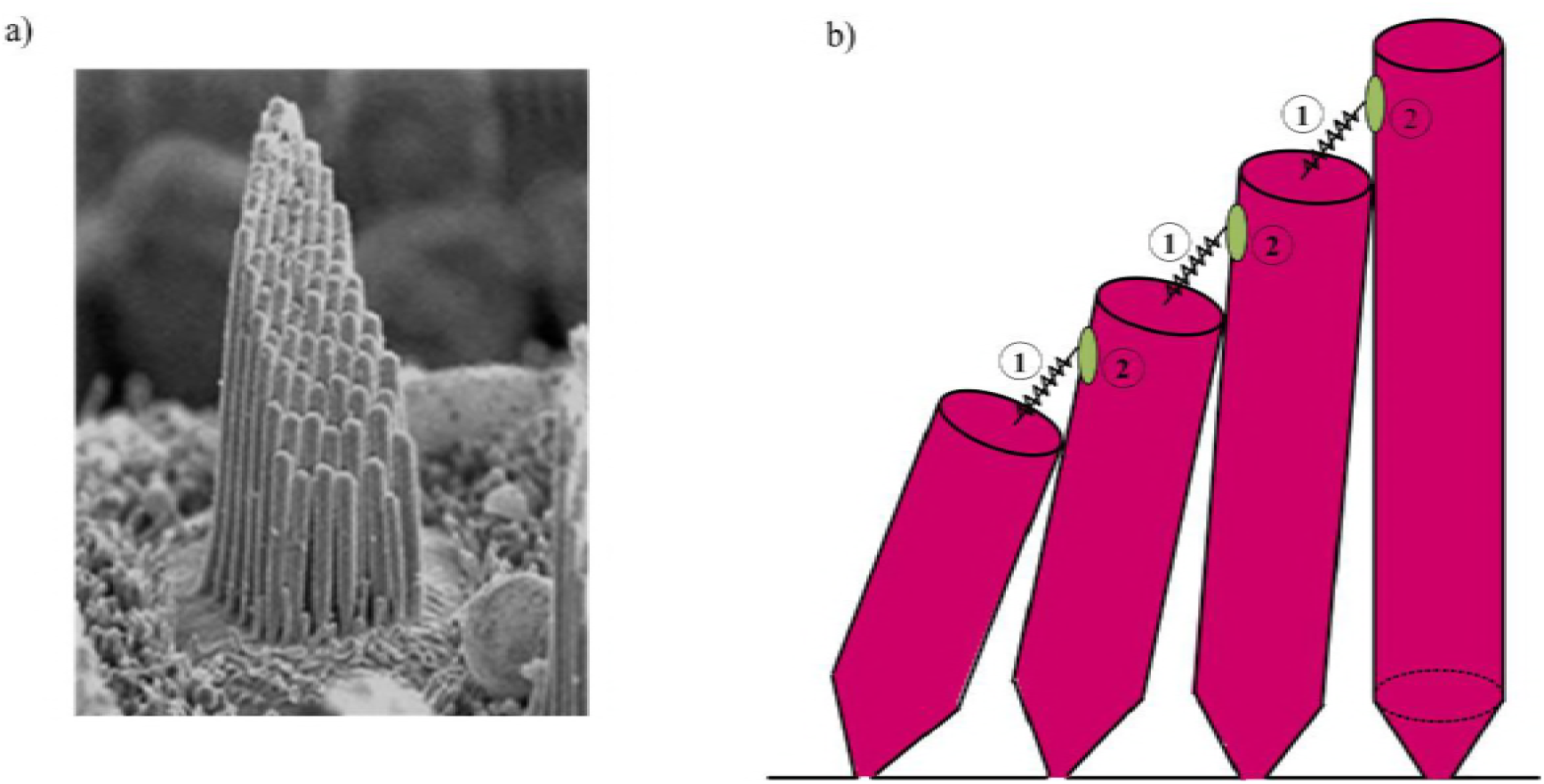
(a) Morphology of a hair–bundle of a vestibular hair cell. In this organ, each bundle consists of approximately 50 stereocilia arranged into several rows with variable heights. (b) Schematic illustrating a single row of stereocilia with tip links–1 and pertaining transduction channels–2, which are responsible for ionic currents.

In response to an external force, a hair–bundle pivots at its base. Let us consider a free standing hair–bundle with highest stereocilia of length 𝓛 and with *b* being the distance between roots of the nearest stereocilia (see Fig. 5a). The position where the tallest stereocilium touches its neighbor is the point O. The distance between the tallest channel and attached motors relative to O is *l*_0_ + *x*_M_, where *l*_0_ is the length of a tip link without extension and *x*_M_ represents the extension under the action of myosin motors. If the tip of the bundle is laterally displaced with distance X (Fig. 5a), all stereocilia pivot at their bases and neighboring ones are sheared by the displacement 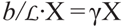. The new location of the touching point between the sterecilia is displaced to O’ making the distance between O’ and the same channel to be *l*_0_ + *x*. Thus, the sheared distance O–O’ is

**Fig. 5.**
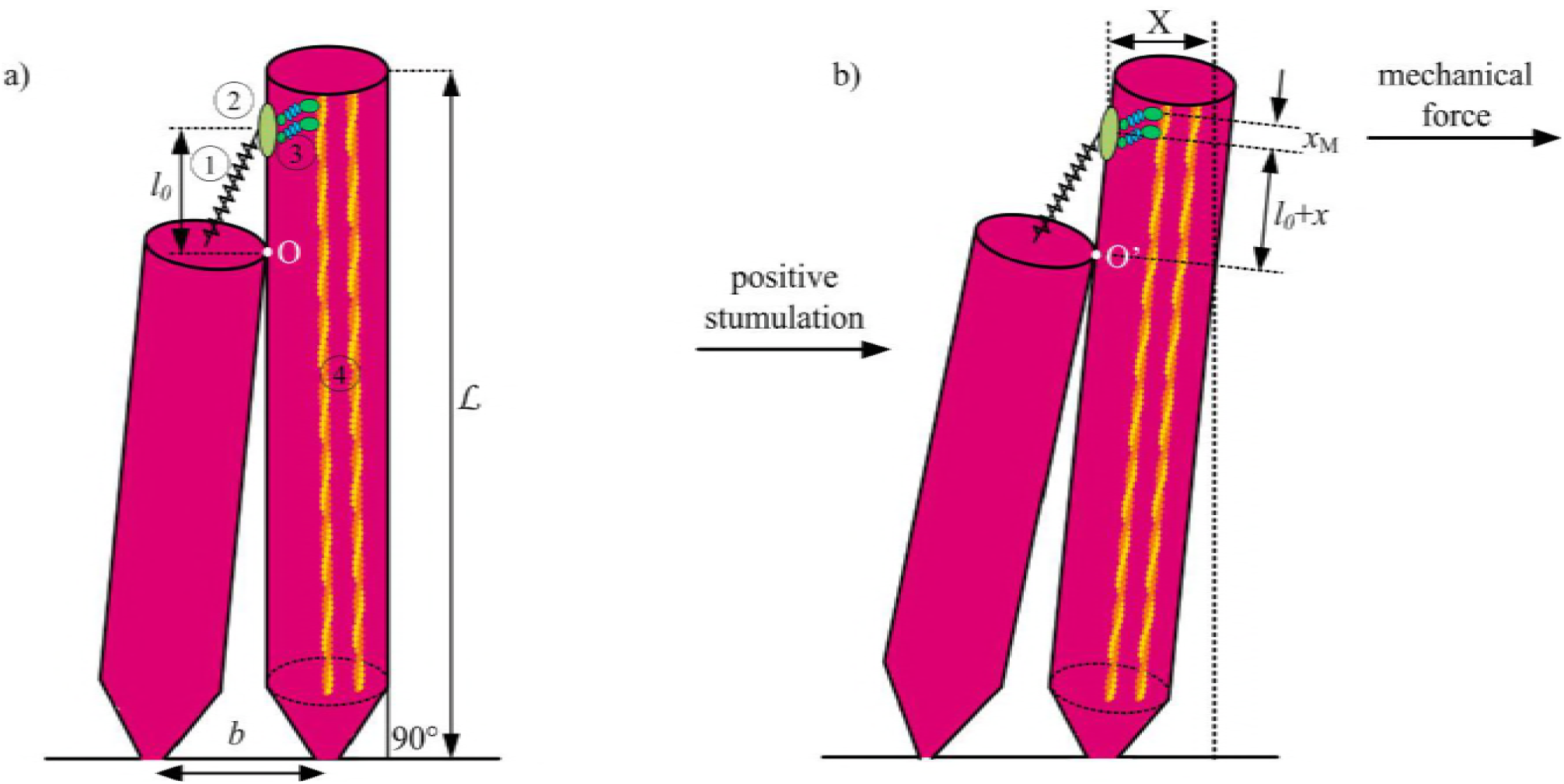
a) The tallest stereocilia of a hair–bundle in equilibrium (1–tip link; 2–transduction channel; 3– myosin adaptation motors; 4–actin filaments). 𝓛 is the height of the tallest stereocilia, b is the distance between the roots of neighboring stereocilia, and *l*_0_ represents the distance between the touching point O and the channel in an equilibrium position of the bundle. b) Transversally displaced bundle by distance X shifts the stereocilia contact point O to new position O^’^ so that its distance is now *l*_0_+*x* with respect to the channel, where *x* stands for channel’s shift due to bundle’s displacement X. This leads to the relation X 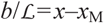.

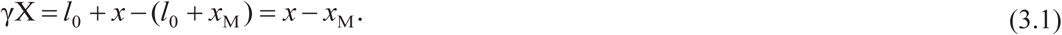

This horizontal displacement of the bundle is countered by the elastic restoring force, which consists of two components. The first one arises from the fact that pivoting around roots stereocilia provides effective bundle stiffness given by

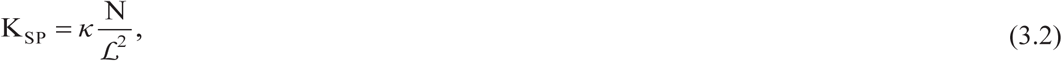

with *κ* being the pivotal stiffness of a single stereocilium. Hudspeth *et al.* [18] revealed the measured value *κ* = 1.5×10^−16^ Nm/rad indicating that the range of K_SP_ is between 0.03 pN/nm for the longest bundles and 11 pN/nm for the shortest ones. The second contribution for the restoring force comes from the tension in the tip links themselves. When the hair–bundle is displaced in positive (right) direction (see Fig. 5b), the gating spring of the tip link exerts a force on the trap door and thus opens the ionic channel. This enables the entry of positive K^+^ and Ca^2+^ ions inside stereocilia. The tip link force consists of two parts in the following form

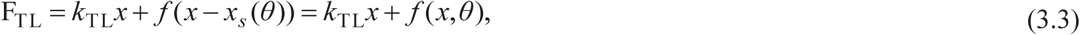

where *x_s_*(*θ*) is an effective shift in the channel gating point dependent on the local Ca^2+^ ion concentration *θ* according to Eq. (1.4). The first term represents the passive linear elastic force with tip link stiffness *k*_TL_, while the second one is the nonlinear active force of transduction channel often called the gating force.

Vilfan and Duke established that the transduction channel protein incorporates a lever arm segment acting as an amplifier for small structural changes as being present when the channel switches state [19]. It turns out that a sufficient model includes just two switching states. In line with that, the open state has the probability *p*_O_(*x*, *θ*) and its corresponding lever arm position is *d*_O_, while the closed state has a probability *p*_C_(*x*, *θ*) and lever arm position *d*_C_. The gating force is related to the channel opening probability of a single open state *p*_O_(*x*, *θ*) by definition [20] whereby

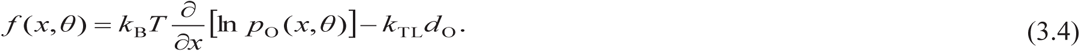

In the steady state the probability for an open state is given in accordance with Boltzmann’s distribution [18]

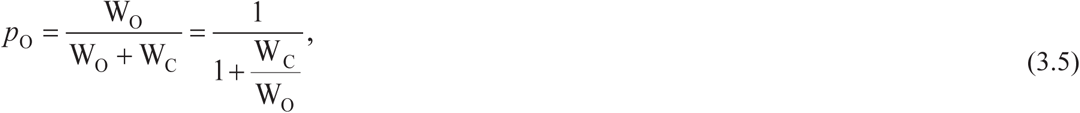

where the weighting functions for open and closed states read:

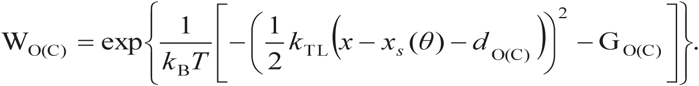

G_O_ and G_C_ are the free energy of open and closed state of transduction channel protein, respectively. They depend on the local Ca^2+^ ions concentration, which is regulated by the entry of Ca^2+^ ions through corresponding channels and by being primarily distributed along actin filaments in the form of mobile “ionic clouds” given by Eq. (2.15).

Based on experimental results [21], the impact of the Ca^2+^ ion concentration on the active force *f*(*x*, *θ*) can be accounted for by an effective shift in the channel gating point, which is related to the local Ca^2+^ ion concentration actually attributed to an incoming “ionic pulse” imparted by the channel’s opening. The shift *x_s_*(*θ*) in the context of our proposed model, see Eq. (2.7), is found as

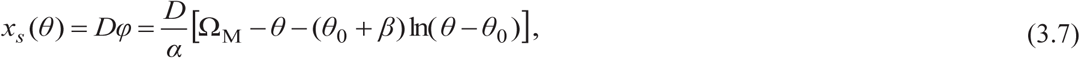

where the constant *D* was empirically estimated to be *D* ~ 4 nm in the bullfrog’s saculus [21]. On the basis of the above, given Eqs. (3.4), (3.5) and (3.6), the explicit form of the gating force for a single channel is

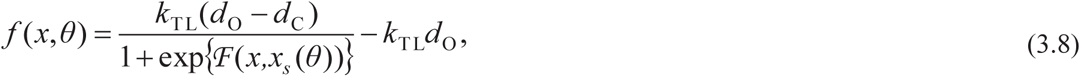

where

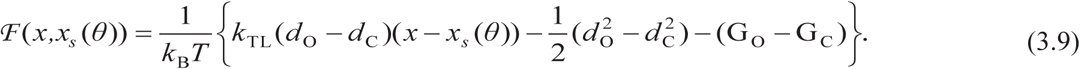

If we use the following set of parameters from the data available in the literature [22, 23]

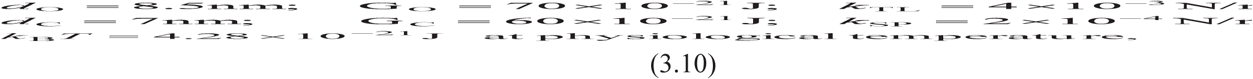

we obtain

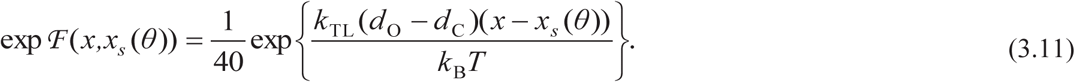

Changing (*x – x_s_*(*θ*)) from zero to infinity, *f*(*x*, *θ*) changes in the narrow interval (– *k*_TL_*d*_C_, – *k*_TL_*d*_O_). Thus we can take the average gating force to be

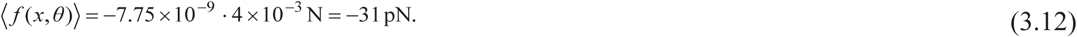

It is also necessary to know the gradient of the gating force, which can be determined from Eqs. (3.8) and (3.9)

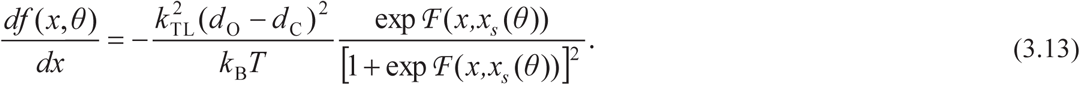

In the similar way as above, and on the basis of the set of parameters given by Eq. (3.10), we estimate the maximum value of Eq. (3.13) taking the maximal value of the exponential factor that amounts to

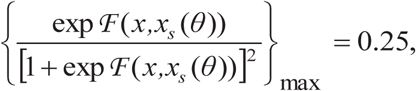

We thus obtain

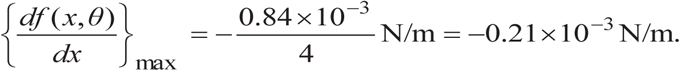

Eventually, the movement of the displaced bundle is also opposed by the viscous drag of the surrounding fluid. The corresponding friction coefficient for bundles of maximal height of sterecilia 𝓛 is

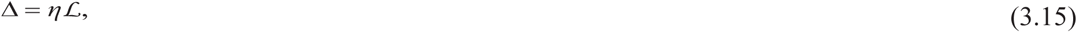

where *η* is the endolymph viscosity.

We stress that the inertial force 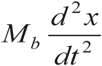, where *M_b_* is the bundle’s mass, is much smaller than the force of viscous damping 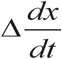 and thus can be safely discarded in the Newtonian equation of motion. By combining Eqs. (3.1), (3.2) and (3.3) including the damping term, the equation of motion for the transduction channel reads

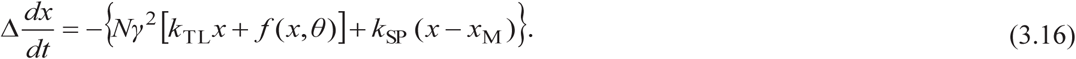

This equation will be exploited in the next section as the basis for establishing a relation between the dynamics of transduction channels and adaptation motors in the stereocilia of a hair–bundle. Eventually, we can make the estimate of the damping parameter Δ. Vilfan and Duke used the value Δ = 0.65 ×10^−6^ Ns/m, which is based on the dynamical viscosity of water [19]. This is much underestimated and we believe that this value should be multiplied by the factor 20 proposed by Dhont and Kang [14]. Consequently, we will use Δ = 13 ×10^−6^ Ns/m in the following calculation.

## 5. The coupled dynamics of adaptation motors and transduction channels of stereocilia

Experimental results [24] revealed that when a vertebrate hair–bundle is suddenly twitched, it exhibits a slow adaptation of the channel current on the order of tens of milliseconds. The transduction channel is linked to the actin filaments of a stereocilium by a number of adaptation motors of myosin– 1c family. It was strongly confirmed that these myosin motors maintain the adequate tension in the tip link thus mediating slow adaptation of a hair bundle [25].

We start from a linear force–velocity relation for these adaptation motors [19]

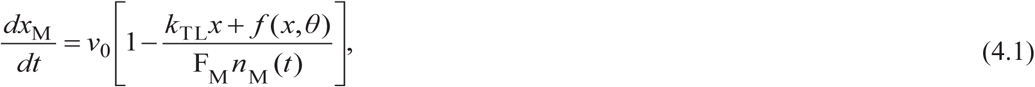

where *x*_M_ is the position of adaptation motors along actin filaments, see Eq. (3.1), *v*_0_ is the zero–load motor velocity, *x* represents the displacement of pertaining transduction channel, see Eq. (3.1), *f*(*x*, *θ*) is the gating force according to Eq. (3.8), F_M_ is the stall force of a single motor and *n*_M_(*t*) is the number of attached adaptation motors.

In order to solve Eq. (4.1), it is first necessary to consider the dynamics of motors detaching from an actin filament governed by the influence of Ca^2+^ ions. Adamek *et al*. [26] showed that Ca^2+^ ions cause a 7–fold inhibition of ATP hydrolysis within a myosin–1c and inversely a 10–fold acceleration of ADP release from it. These effects lead to the increase of the detachment rate of the motor–filament cross–bridge and increase the lifetime of the motors’ detached state. The detachment dynamics *n*_M_(*t*) of initially present *N*_M_ = 20 motors bound to an actin filament [19] is governed by the following linear equation

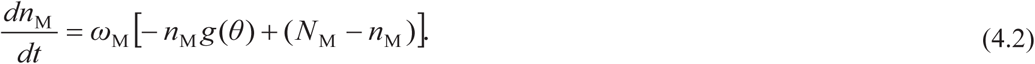

Here, *ω*_M_ = 200 s^−1^ stands for the myosin binding rate, while *g*(*θ*) is a nonlinear function describing how the concentration of Ca^2+^ ions impacts the motor’s detachment rate. Following the method used by Vilfan and Duke [19] we assume that *g*(*θ*) has the cubic nonlinear form with respect to the dimensionless excess concentration *θ* of Ca^2+^ ions in the mobile “ionic pulse” around actin filaments

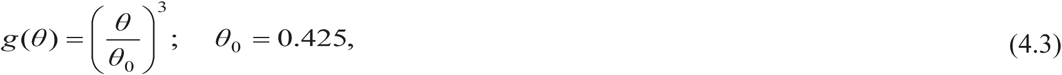

Then, Eq. (4.2) can be solved exactly and we obtain the number of remaining attached motors as the function of time, with the initial condition *n*_M_(*t*=0) = *N*_M_,

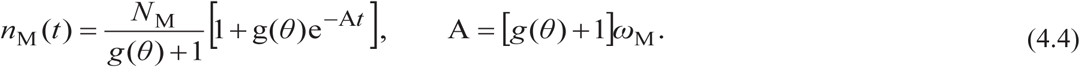

The characteristic detachment time *τ_d_* is represented in Fig. 6 and it is given by

**Fig. 6.** Characteristic detachment time *τ_d_* for *θ* = 0.45 (left) and *θ* = 0.95 (right).

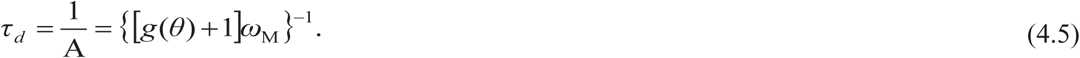

Since within the different ionic pulses the concentration *θ* could change in the range *θ*_0_ < *θ* ≤ 2.35*θ*_0_, it implies that the following inequality is valid

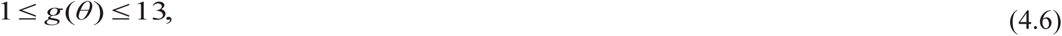

and the range of the detachment rate is, respectively

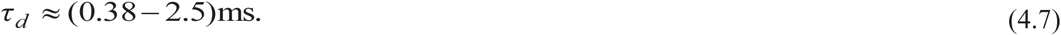

This is in sharp contrast with the statements given by Vilfan and Duke [19] in which they pointed out that the motors operate on a slower timescale than mechanical relaxation of a bundle. Hence, in their description of bundle’s dynamics the motor position *x*_M_ was taken as being fixed.

This new assumption allows us to safely link the dynamics of adaptation motors given by Eq. (4.1) with the Eq. (3.16) that describes the motion of the transduction channels. In order to combine these equations, we first differentiate the channel’s equation with respect to time as

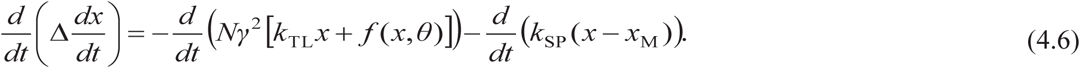

Then using Eq. (4.1), we get the following second order differential equation with respect to *x*(*t*)

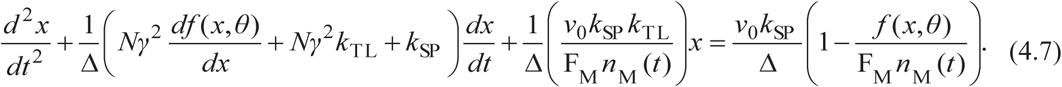

Strictly speaking, this equation is nonlinear due to the presence of the exponential function *f*(*x*, *θ*) and *df*(*x*, *θ*)/*dx* as well as function *n*_M_(*t*). However, using the previously found numerical estimates of the narrow ranges of these functions given in Eqs. (3.12) and (3.14), nonlinearity of above equation is eliminated. Also, it is suitable, instead of the explicit function *n*_M_(*t*) defined by Eq. (4.4), to keep here just a characteristic numbers of motors remaining after the detachment time *τ_d_*, namely *n*_M_ = *N*_M_/14 = 10/7 for the largest influx giving the fastest “ionic pulse” arising in the open channel state, and *n*_M_ = *N*_M_/2 = 10 for the smallest influx of Ca^2+^ ions and the slowest “ionic pulse”.

Furthermore, we can use the following abbreviations in Eq. (4.7)

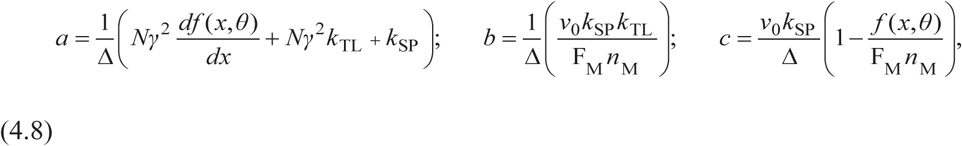

so that the compact form of the equation of motion now reads

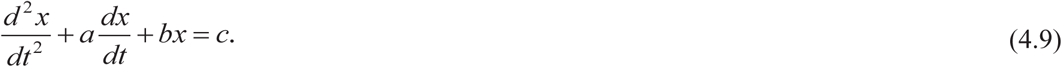

The solution of the homogeneous part of the above equation is

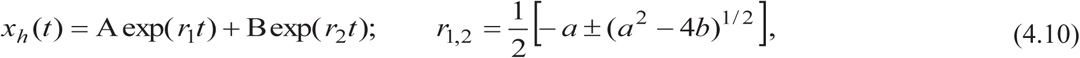

where the integration constants A and B are defined by the initial conditions.

Of particular interest here is to determine coefficients *a*, *b* and *c* defined by Eq. (4.8). Using the set of estimated parameters given by Eqs. (3.10), (3.12), (3.14), and (3.17) completed with F_M_ = 1.25 pN, *v*_0_ = 0.3 μm/s, *N*_M_ = 20, and *n*_M_ = 10/7 [19], the following values are obtained

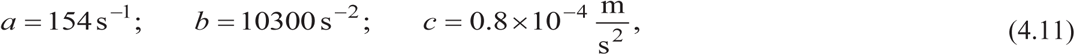

which lead to the explicit parameters 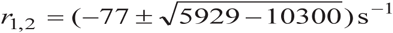. This describes the damped oscillations with the decay rate

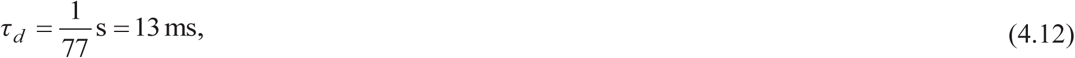

which is in accordance with the experiment of Wu *et al.* [24]. Also, this value is one order of magnitude greater than the detachment time of adaptation motors, Eq. (4.7). The corresponding angular velocity of these relaxation oscillations is *ω* = (4371)^1/2^s^−1^ = 66.1 rad/s, giving the frequency

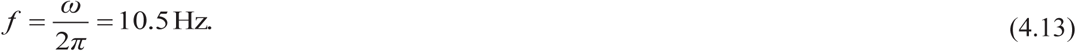

This value is in good agreement with the available experimental results [23, 27]. Tinevez *et al.* [27] demonstrated that a hair–bundle from bullfrog saculus exhibits spontaneous oscillations at a frequency of ~ 12 Hz and a round–mean–squared magnitude of 22 nm.

Finally, a particular solution of Eq. (4.9) is found as

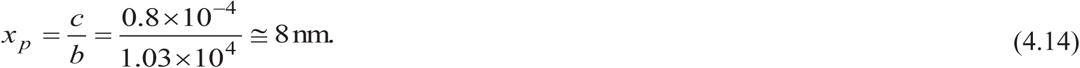

This value is very close to *d* = 7 nm, which is the distance of gating spring relaxation on channel opening [22]. From Eq. (4.1) it follows that the condition for the emergence of the bundle’s spontaneous oscillations is given by

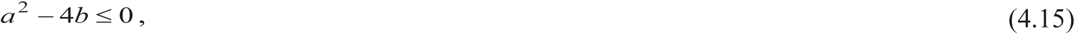

which yields

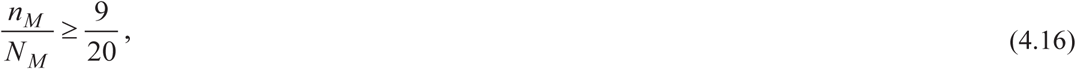

meaning that the number of attached motors must be above 45% in order to self–tune such an active process. Increasing the Ca^2+^ ion concentration *θ* increases the value of parameter *b*, which leads to an increase in the frequency of oscillations. Sinusoidal oscillations defined by Eq. (4.10) that are specified by parameters given in Eqs. (4.12) and (4.13) correspond to the leading harmonic. The higher harmonics reflecting the real experimental bundle movement in bullfrog sacculus [23] manifest the basically nonlinear character of Eq. (4.7) due to the functions *f*(*x*, *θ*), *df*(*x*, *θ*)/*dx* and *n*_M_(*t*). These harmonics have not been considered here. All these arguments clearly corroborate the validity of our concept in establishing a strong correlation between the dynamics of transduction channels and active movements of myosin adaptation motors in hair–bundles mediated by Ca^2+^ ions carried by propagating localized “ionic pulses” described by our model in Eq. (2.7).

## 6. Discussion and conclusions

Experimental and theoretical studies have shown that the non–mammalian hair–bundles can produce a force due to the action of Ca^2+^ ions. This active force production is connected to the fast adaptation of the transduction Ca^2+^ currents. The main interest of this paper is centered on the specific mechanism of propagation of Ca^2+^ ions from transduction channels of hair cells of the non– mammalian vertebrate inner ear apparatus. The polyelectrolyte properties of actin filaments explain the reasons why Ca^2+^ ions form dense “ionic clouds” around these protein filaments with local concentrations of these ions much higher than in the bulk solution. These Ca^2+^ ions propagate along actin filaments with a velocity proportional to the density of the “ionic cloud” and substantially greater than the diffusion velocity of free ions. This concept leads to a mechanism with the detachment rate of adaptation myosin motors, which is faster, in the range of milliseconds as given in Eq. (4.7), than that obtained in the context of the theory established by Vilfan and Duke [19]. Consequently, we have been able to unify the dynamics of adaptation myosin motors and transduction channels. This leads to damped oscillations of the transduction channels according to Eqs. (4.10), (4.11) and (4.13). The relaxation rate of these oscillations is on the order of ten milliseconds, which is one order of magnitude longer than the motor’s detachment time. This fact justifies our replacement of the explicit function *n*_M_(*t*) defined by Eq. (4.4) with a characteristic number of motors remaining after rapid detachment caused by a fast sweeping movement of the “ionic cloud”. We stress that the frequency of the above oscillations is the order of ten Hertz, which is very close to the available experimental data on the spontaneous oscillations of a hair–bundle in the bullfrog saculus [23, 27].

## Acknowledgments

This research was financially supported by the Provincial Secretariat for Higher Education and Scientific Research of AP Vojvodina (Project No. 114–451–2708/2016–03), also by the Ministry of Education, Science and Technological Development of the Republic of Serbia (Projects No. OI171009 and III43008), and by the Serbian Academy of Sciences and Arts.

## Appendix 1

We consider the case that the influx of Ca^2+^ ions provides a locally elevated concentration *θ*(*x*,*t*) > *θ*_0_, which flows along an actin filament in the traveling wave form

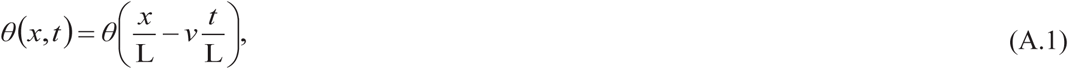

where L = 5.4 nm stands for the length of the filament’s unit and *v ≤ v*_0_ is the wave velocity. In order to make the master equation dimensionless, it is convenient to introduce new variables and parameters as follows

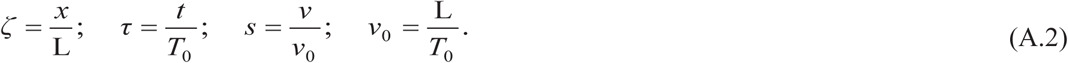

By analogy with solitonic waves, we postulate that the velocity *v* of an “ionic pulse” is proportional to its density, namely the number of involved Ca^2+^ ions, which is provided by the intensity of the permitted ionic influx through the corresponding transduction channel and expressed by *θ*_M_, the maximum of the relative concentration of Ca^2+^ ions within an “ionic pulse”. Furthermore, the form of a traveling wave enables that time and space can be unified with a single dimensionless variable *φ*

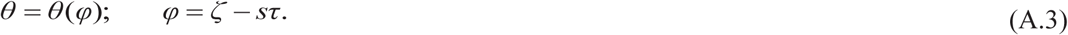

After performing the straightforward transformation as defined above, the partial differential equation given by Eq. (2.6) becomes an ordinary differential equation of the following form

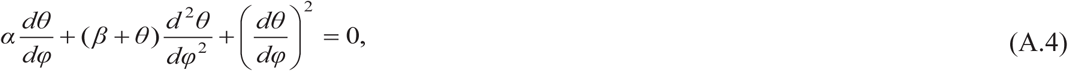

where two abbreviations were introduced, namely

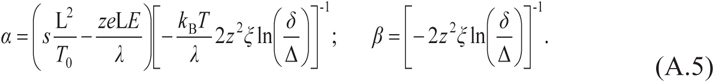

The order of Eq. (A.4) can be reduced by the standard substitution:

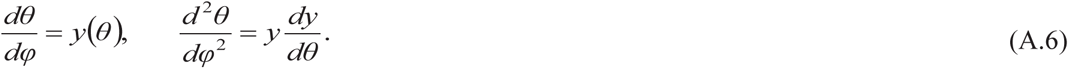

It leads to the linear differential equation of first order in the following form

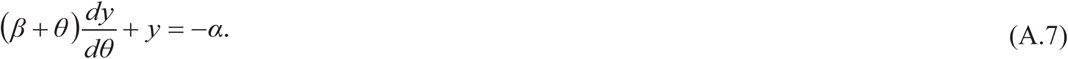

The next step is to solve this equation in terms of the following improved boundary conditions in comparison with our previous calculation [1]

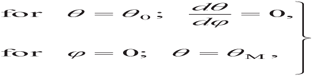

where *θ*_M_ represents the local maximum of concentration of injected Ca^2+^ ions within the “ionic pulse”, while *θ*_0_ has the meaning in accordance with Eq. (1.11). After a straightforward calculation and finding the constants of integration from the above conditions, the solution of Eq. (A.7) in the form of an implicit function *θ*(*φ*) is

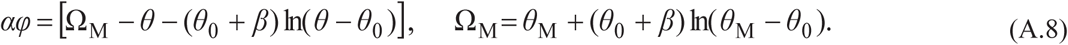

## References

[1] Sataric, M.V., Sekulic, D.L., Sataric, B.M.: Actin filaments as the fast pathways for calcium ions involved in auditory processes. J. Biosciences 40, 549–559 (2015)

[2] Manning, G.S.: Approximate solutions to some problems in polyelectrolyte theory involving nonuniform charge distributions. Macromolecules 41, 6217–6227 (2008)

[3] Tang, J.X., Janmey, P.A.: The polyelectrolyte nature of F–actin and the mechanism of actin bundle formation. J. Biol. Chem. 271, 8556–8563 (1996)

[4] Gartzke, J., Lange, K.: Cellular target of weak magnetic fields: ionic conduction along actin filaments of microvilli. Am. J. Physiol. Cell Physiol. 283, C1333–C1346 (2002)

[5] Sataric, M.V., Bednar, N., Sataric, B.M., Stojanovic, G.: Actin filaments as nonlinear RLC transmission lines. Int. J. Mod. Phys. B 23, 4697–4711 (2009)

[6] Manning, G.S.: A counterion condensation theory for the relaxation, rise, and frequency dependence of the parallel polarization of rodlike polyelectrolytes. Eur. Phys. J. E 34, 39 (2011)

[7] Israelachvili, J.N.: Intermolecular and surface forces: With applications to colloidal and biological systems. Academic Press, London (1992)

[8] Beurg, M., Nam, J.H., Chen, Q., Fettiplace, R.: Calcium balance and mechanotransduction in rat cochlear hair cells. J. Neurophysiol. 104, 18–34 (2010)

[9] Lin, E.C., Cantiello, H.F.: A novel method to study the electrodynamic behavior of actin filaments. Evidence for cable–like properties of actin. Biophys. J. 65, 1371–1378 (1993)

[10] Minoura, I., Muto, E.: Dielectric measurement of individual microtubules using the electroorientation method. Biophys. J. 90, 3739–3748 (2006)

[11] Uppalapati, M., Huang, Y.M., Jackson, T.N., Hancock, W.O.: Microtubule alignment and manipulation using AC electrokinetics. Small 4, 1371–1381 (2008)

[12] Cantero, M.R., Perez, P.L., Smoler, M., Etchegoyen, C.V., Cantiello, H.F.: Electrical oscillations in two–dimensional microtubular structures. Sci. Rep. 6, 27143 (2016)

[13. Sekulic, D.L., Sataric, B.M., Zdravkovic, S., Bugay, A.N., Sataric, M.V.: Nonlinear dynamics of C–terminal tails in cellular microtubules. Chaos 26, 073119 (2016)

[14] Dhont, J.K.G., Kang, K.: Electric–field–induced polarization and interactions of uncharged colloids in salt solutions. Eur. Phys. J. E 33, 51–68 (2010)

[15] Ashmore, J., Avan, P., Brownell, W.E., Dallos, P., Dierkes, K., Fettiplace, R., Grosh, K., Hackney, C.M., Hudspeth, A.J., Jülicher, F., Lindner, B., Martin, P., Meaud, J., Petit, C., Santos Sacchi, J.R., Canlon, B.: The remarkable cochlear amplifier. Hearing Res. 266, 1–17 (2010)

[16] Benser, M.E., Issa, N.P., Hudspeth, A.J.: Hair–bundle stiffness dominates the elastic reactance to otolithic–membrane shear. Hear. Res. 68, 243–252 (1993)

[17] Holt, J.R., Corey, D.P.: Two mechanisms for transducer adaptation in vertebrate hair cells. Proc. Natl. Acad. Sci. USA 97, 11730–11735 (2000)

[18] Hudspeth, A.J., Choe, Y., Mehta, A.D., Martin, P.: Putting ion channels to work: Mechanoelectrical transduction, adaptation, and amplification by hair cells. Proc. Natl. Acad. Sci. USA 97, 11765–11772 (2000)

[19] Vilfan, A., Duke, T.: Two adaptation processes in auditory hair cells together can provide an active amplifier. Biophys. J. 85, 191–203 (2003)

[20] van Netten, S.M., Kros, C.J.: Gating energies and forces of the mammalian hair cell transducer channel and related hair bundle mechanics. Proc. R. Soc. Lond. B 267, 1915–1923 (2000)

[21] Corey, D.P., Hudspeth, A.J.: Kinetics of the receptor current in bullfrog saccular hair cells. J. Neurosci. 3, 962–976 (1983)

[22] Martin, P., Mehta, A.D., Hudspeth, A.J.: Negative hair–bundle stiffness betrays a mechanism for mechanical amplification by the hair cell. Proc. Natl. Acad. Sci. USA 97, 12026–12031 (2000)

[23] Martin, P., Bozovic, D., Choe, Y., Hudspeth, A.J.: Spontaneous oscillation by hair bundles of the bullfrog's sacculus. J. Neurosci. 23, 4533–4548 (2003)

[24] Wu, Y.C., Ricci, A.J., Fettiplace, R.: Two components of transducer adaptation in auditory hair cells. J. Neurophysiol. 82, 2171–2181 (1999)

[25] Holt, J.R., Gillespie, S.K.H., Provance, D.W., Shah, K., Shokat, K.M., Corey, D.P., Mercer, J.A., Gillespie, P.G.: A chemical–genetic strategy implicates myosin–1c in adaptation by hair cells. Cell 108, 371–381 (2002)

[26] Adamek, N., Coluccio, L.M., Geeves, M.A.: Calcium sensitivity of the cross–bridge cycle of Myo1c, the adaptation motor in the inner ear. Proc. Natl. Acad. Sci. USA 105, 5710–5715 (2008)

[27] Tinevez, J.Y., Jülicher, F., Martin, P.: Unifying the various incarnations of active hair– bundle motility by the vertebrate hair cell. Biophys J. 93, 4053–4067 (2007)

